# Elevating Neuronal *CYLD* Causes Frontotemporal Dementia (FTD)-Relevant Behavioral and Physiological Deficits

**DOI:** 10.64898/2026.02.24.707507

**Authors:** Aparajita Baral, Mohammad Bilal, Huihui Dai, Yong-Woo Jun, Sandra Almeida, Fen-Biao Gao, Wei-Dong Yao

**Affiliations:** Departments of Psychiatry & Behavioral Sciences, and of Neuroscience and Physiology, State University of New York, Upstate Medical University, Syracuse, NY 13210, USA; Frontotemporal Dementia Research Center, RNA Therapeutics Institute, University of Massachusetts Chan Medical School, Worcester, MA 01605, USA; Department of Neurology, University of Massachusetts Chan Medical School, Worcester, MA 01605, USA

## Abstract

Frontotemporal dementia (FTD), a leading form of presenile dementia disrupting behavior, language and/or movement, is linked to mutations of a number of genes, including *CYLD* that encodes a Lys63 (K63) deubiquitinating enzyme. Among several *CYLD* variants found in FTD patients, a gain-of-function missense mutant, M719V, has been proposed to be pathogenic, but its pathogenicity in vivo and the underlying mechanism remain unknown. Here, we have developed transgenic mice that express either wildtype (WT) or M719V-CYLD in neurons throughout the mouse brain using adeno-associated virus (AAV) mediated somatic brain transgenesis. We show that somatic *M719V-CYLD* transgenic mice display profound FTD-associated behavioral impairments, including risk-taking, reduced social interaction, and loss of empathy that emerge from early stages and worsen with aging. Furthermore, *M719V-CYLD* mice also show significant early neurophysiological impairments in the prefrontal cortex (PFC), including depolarized resting membrane potential, decreased synaptic transmission, and reduced neuronal excitability. Surprisingly, however, *M719V-CYLD* mouse brain exhibits elevated autophagy activity and decreased Akt-mTOR signaling without overt neuronal cell loss or microgliosis even at 12 months of age. Most *M719V*-*CYLD*-associated cellular and behavioral phenotypes are also recapitulated but to a lesser extent in *WT-CYLD* mice, suggesting *CYLD* activation is responsible for the observed neural circuit deficits and the M719V mutation is gain-of-function in nature. Our results uncover important roles of neuronal CYLD in PFC function and social behaviors and establish a unique animal model to investigate pathogenic mechanisms of FTD, in particular its social behavioral deficits, at molecular, cellular, synaptic and circuit levels.

## Introduction

Frontotemporal dementia (FTD) is caused by focal and progressive atrophy of frontal and/or anterior temporal cortices and is the leading form of dementia under the age of 60 [1, 2]. Predominant symptoms of FTD are behavioral abnormalities characterized by changes in personality, loss of empathy, apathy, disinhibition, and/or language deficits at early-middle stages, and general memory and cognitive deteriorations at later stages [1, 2]. Thus, early prodromal synaptic and circuit dysfunctions mediating behavior impairments may precede massive neurodegeneration and disability. Cardinal pathological features of FTD include nuclear depletion and cytoplasmic inclusions of TDP-43 (transactive response-DNA binding protein-43), astrogliosis, microgliosis, and neuronal cell loss [2, 3]. About 40% of FTD are familial, associated with mutations of at least 15 genes of diverse molecular functions [2, 4, 5]. Remarkably, at least 10 of these genes are involved in autophagy, a conserved cell quality-control process that clears damaged or unwanted cytoplasmic content through lysosomal degradation [6, 7], suggesting a central role of autophagy dysregulation in FTD pathogensis [4, 8, 9]. FTD is linked clinically, pathologically, genetically, and mechanistically to the motor neuron disease amyotrophic lateral sclerosis (ALS) [2, 10]. FTD pathogenesis also has significant implications for Altheimer’s disease (AD) [11].

A recently identified FTD risk gene is *CYLD*, which encodes a Lys63 (K63) deubiquitinating enzyme (DUB) [12]. A missense mutation of *CYLD* (*M719V*) was first discovered in a large European Australian family with autosomal dominant inheritance of FTD/ALS [3], and shortly after, two other variants have been reported in a Portuguese cohort [13] followed by additional rare variants found in Chinese ALS patients [14] and other disease-associated mutations [15]. Interestingly, CYLD interacts with products of three FTD genes *p62/SQSTM1* (autophagy receptor), *Optineurin* (autophagy receptor), and *TBK1* (a serine/threonine kinase that phosphorylates and activates p62 and Optineurin) [16–21], suggesting a potential role for CYLD in autophagy related to FTD. M719V-CYLD mutant exhibits higher K63-deubiquitase (DUB) enzyme activity and patients show inclusions of TDP-43, the major pathologic component of tau-negative and ubiquitin-positive inclusions in ∼50% of FTD patients [22–24] that is cleared by autophagy [25, 26]. How M719V-CYLD mutant drives FTD pathogenesis in animal models or human neurons are unknown.

CYLD is best known as a negative regulator of NF-κB activation in immune signaling [27] and was initially identified as a tumor suppressor whose loss of function mutations are linked to familial cylindromatosis [28–30]. More recently, however, we and others have shown that CYLD is present in excitatory neurons and enriched in the postsynaptic density (PSD) [31], trafficks to PSD in response to neuronal activity and regulates PSD assembly [32–34], interacts with numerous disease-related synaptic proteins [35], regulates dendritic spine formation and maintenance and synaptic plasticity [31, 36], and modulates neuronal excitability [37] and behaviors [36, 38, 39]. Importantly, we have shown that CYLD stimulates the induction of neuronal autophagy by inhibiting Akt-mTOR signaling, driving synapse pruning and inhibiting synaptic plasticity [36]. How FTD-associated *CYLD* mutations impair neuronal excitability and synaptic efficacy have not been examined.

Here, we develop AAV-mediated somatic transgenic mouse models that express M719V-CYLD in postnatal neurons of the mouse brain. We show that *M719V-CYLD* somatic transgenic mice develop FTD-associated behavioral and neurophysiological impairments that emerge early and worsen with aging, but without overt FTD-related cellular pathologies such as neuronal cell loss. Those phenotypes are also recapitulated but to a lesser extent in *WT-CYLD* mice. Our results reveal important roles of neuronal CYLD in the maintenance of prefrontal circuit functions important for FTD-disrupted behaviors and establish a unique animal model to investigate early prodromal pathogenic mechanisms of *CYLD*-FTD at molecular, cellular, synaptic and circuit levels.

## Materials and Methods

### Animals

C57BL/6J mice were housed 2-5 per cage under a 14/10-hour light/dark cycle with *ad litbitum* access to standard chow food and water. Both male and female mice were used, and data was combined. The housing environment in the animal facility was maintained at 21-23 °C and 40-70% humidity. All studies were conducted in accordance with the guidelines of the National Institutes of Health Guide for the Care and Use of Laboratory Animals approved by the SUNY Upstate Medical University institutional animal care and use committees.

### Viral constructs and packaging

To generate pAAV-Syn-3xHA-CYLD wild-type or M719V constructs, the coding sequence of human CYLD wild-type (NM_001378743.1) or the M719V mutant fused to an N-terminal 3×HA tag was synthesized by Genewiz and cloned into the pAAV-Syn1-MCS vector (a gift of Dr. Yingxi Lin) using NheI and AscI restriction sites. To generate pAAV-Syn-EGFP-CYLD wild-type or M719V constructs, EGFP-CYLD wild-type or M719V fragments were amplified by PCR using EGFP-NotI-a-S (5’-ATAAGAATGCGGCCGCaGCCACCATGGTGAGCAAGGG-3’) and CYLD-SalI-A (5’-CACGCGTCGACttatttgtacaaactcattg-3’), and the PCR products were digested with NotI and SalI and subcloned into the corresponding sites of the *pAAV-hSyn-EGFP* plasmid, which was kindly provided by The Viral Vector Core (VVC) of the Gene Therapy Center at UMass Chan Medical School. CYLD and mutant constructs were verified with automated sequencing. High titer AAVs were packaged at Boston Children’s Hospital Viral Core.

### Surgeries

Generation of somatic transgenic mice with neonatal intracerebroventricular (ICV) viral injections have been described previously [40, 41]. Briefly, newly born pups on postnatal day 0 (p0) were cryoanesthetized until there was no movement. ICV microinjections of high titer (1.2-2 x 10^13^) AAVs were achieved with a 32-gauge needle attached to a 10 mL syringe (Hamilton Company). Virus injection groups included: AAV2/9-hSyn-EGFP, AAV2/9-hSyn-EGFP-CYLD, AAV2/9-hSyn-EGFP-M719V-CYLD, AAV2/9-hSyn-EGFP and AAV2/9-hSyn-3xHA-CYLD, and AAV2/9-hSyn-EGFP and AAV2/9-hSyn-3xHA-M719V-CYLD. Following injections, the pups were placed on a heating pad to recover before moving back into their home cages.

Stereotaxic injections of AAVs into the mPFC have been detailed previously [41]. Mice were anesthetized with 1.5%–2% isoflurane and placed on a stereotaxic apparatus (Kopf Instruments) under aseptic conditions. Burr holes were made just above the injection sites on both hemispheres using a drill. Bilateral viral injections of 0.25 µL were made using a Neuros syringe (Hamilton Company) at a rate of 0.05 µL per minute with a micromanipulator (Stoelting Company) into the prelimbic area. The coordinates were anterior-posterior (AP) +1.94 mm, mediolateral (ML) ±0.350 mm, and dorsoventral (DV) −1.90 mm (from Bregma). After completing the injections, the needles were left in place for 10 minutes to ensure complete delivery of the virus, and then slowly withdrawn with caution.

### Behaviors

Open Field Test (OFT) [36]: Mice were placed in the center of an open field arena (Med Associates) and allowed to explore for 1 hour freely. Mouse behaviors were recorded by Any maze Software (ANY-Maze, Stoelting). Parameters measured included the total distance traveled, the time spent in the center zone and edge zone of the apparatus. And the ratio of the two.

Elevated Plus Maze (EPM) [36]: The elevated plus maze consisted of an open arm and a closed arm. A test mouse was placed in the interaction area at the beginning and allowed to explore all arms for 5 minutes freely. The total time spent on each arm and on the edge of the open arm was scored.

Three chamber social behavioral test [36]: A classical 5-session assay was used. A test mouse was placed in the center chamber for 5 minutes. After 5 minutes, dividers were removed on either side, and the mouse was allowed to move freely to explore all three chambers for 10 minutes. On day 1, a stranger mouse of the same sex, age, and strain was placed in an inverted cup in the corner of the east chamber. On days 2-4, the same mouse was placed back in the same inverted cup for the tests. On day 5, a novel stranger mouse of the same gender and age was placed in the inverted empty cup in the west chamber and the test mouse was allowed to interact freely with the novel and the now familiar mice. The total time spent interacting with each mouse during all 5 days for sociability and social memory and time spent in each chamber were measured.

Distress-induced affiliative (DIA) assay [41]: All mice were handled and habituated before test. On test day, an unfamiliar mouse (demonstrator) of the same gender and age was placed in the homecage of an observer mouse (observer) and allowed to freely interact for 15 minutes (Homecage 1 or HC1). Immediately after HC1, the demonstrator mouse was placed in a fear conditioning chamber and received repeated electric shocks on the feet, with the observer mouse was placed in an adjacent chamber freely observing. This 9-min observational fear conditioning (OFC) session consisted of 5 min of habituation followed by 4 minutes of foot shocks (2 ms shock every 10 s, 1 mA). After the OFC, the demonstrator was placed back to the observer’s home cage and allowed to freely interact for 15 min (HC2). Throughout the sessions, behaviors of both observer and demonstrator mice were recorded with a ceiling-mounted camera and scored with Any-Maze or manually. The following parameters were measured: allogrooming and body contact affiliative behavior in observers toward demonstrators, social investigation behavior (sniffing), and non-social behavior (rearing and self-grooming) during HC1 and HC2 and observer freezing response in OFC.

### Immunoblotting

Cortical tissues were lysed and protein concentrations determined using a Bradford protein assay. For Western blotting, samples were run on SDS-PAGE gels to separate proteins, and the blots were transferred to PVDF membranes. The membranes were incubated with primary antibodies (1:1000) against total and phosphorylated AKT and mTOR, p62, and LC3. Appropriate secondary antibodies were used at a 1:2000 dilution. Beta-actin was used as the loading control. The blots were developed, and protein bands were quantified by densitometry within a linear range.

### Immunohistochemistry, confocal microscopy and imaging analysis

Mice were sacrificed by cervical dislocation and brains quickly removed. Brains were fixed in 4% paraformaldehyde prepared in 1× PBS for 24-48 hours at 4 °C. Brains were sectioned at 20 µm using a cryostat, and the sections were collected and stored at 4 °C. Sections were then incubated in permeabilization solution for 2 hours in 3% BSA in 1× PBS at room temperature with primary antibodies - NeuN (anti-mouse), p62 (anti-rabbit), CD11b (anti-rat), and IBA1 (anti-rabbit) - prepared in 3% Triton X-100 in BSA solution and incubated overnight in a cold room. After rinsing three times with 1× PBS, secondary antibodies were added: Alexa Fluor 568 (anti-rabbit) for p62, and IBA1, Alexa Fluor 647 (anti-rat) for CD11b, and Alexa Fluor 568 (anti-mouse) for NeuN, all at a 1:500 dilution. After incubation at room temperature for 2 hours, tissue slices were carefully transferred onto glass slides. ProLong Gold Antifade mounting medium with DAPI was used. Images were acquired using a confocal microscope and analyzed with ImageJ software.

### Electrophysiology

Mice injected (either via ICV or via stereotaxically to the PL) with AAVs encoding EGFP, EGFP-CYLD or EGFP-M719V were sacrificed by cervical dislocation and their brains rapidly removed and placed into ice-cold artificial cerebrospinal fluid (ACSF) containing (in mM): 126 NaCl, 2.5 KCl, 2.5 CaCl_2_, 1.2 MgCl_2_, 25 NaHCO_3_, 1.2 NaH_2_PO_4_ and 11 D-glucose. 300-μm coronal mPFC slices containing the PL were cut in ice-cold oxygenated ACSF (with 95% O_2_ and 5% CO_2_) using a vibratome (Leica). Brain slices were incubated in oxygenated ACSF at 21-23 °C for at least 1 hour before electrophysiological recording.

Whole-cell patch-clamp electrophysiology was performed as previously described [36, 41, 42]. EGFP-positive and negative pyramidal neurons in the prelimbic cortex were visualized under DIC and green fluorescence under an upright microscope (BX51WI, Olympus). For current-clamp recording, pipettes (3.5–4.5 MΩ) were filled with (in mM) 130 K-gluconate, 8 NaCl, 10 HEPES, 0.4 EGTA, 2 Mg-ATP, and 0.25 GTP-Tris, pH 7.25. Intrinsic excitability was assessed by depolarizing step- or ramp current currents in the presence of picrotoxin (50 μM). Input resistance was measured by a series of hyperpolarizing currents of 500 ms, −25 pA current steps at 10 s intersweep intervals. For voltage-clamp recording, pipettes (4.5–5.5MΩ) were filled with (in mM): 142 Cs-gluconate, 8 NaCl, 10 HEPES, 0.4 EGTA, 2.5 2 Mg-ATP, and 0.25 GTP-Tris, QX-314, pH 7.25 (with CsOH). Miniature excitatory synaptic currents (mEPSCs) were recorded in the presence of tetrodotoxin at −70 mV. AMPA/NMDA ratio was measured with a kinetics based method [42], defined as the amplitude of the NMDAR component 80 ms post-stimulation at +40 mV divided by the peak AMPAR component at −70 mV. All recordings were made with a Multiclamp 700B amplifier (Molecular Devices) at 32 °C with a temperature controller (Harvard Apparatus). Data acquisition and analysis were done using Digidata 1322A and pClamp software (version 9.2; Molecular Devices) or MiniAnalysis. Signals were digitized at 20 kHz and filtered off-line at 2 kHz.

### Statistics

Data are shown as mean ± SEM. Statistical analysis was performed using one-way or two-way ANOVA followed by Tukey’s multiple comparison post-hoc test, or Kolmogorov-Smirnov test, as specified in individual figures. All statistical analyses were performed using Prism 10.0 (GraphPad Software).

## Results

### Generating a somatic transgenic mouse model of *CYLD*-FTD

To delineate the pathogenic potential and underlying mechanisms associated with *M719V-CYLD* in vivo, we first sought to establish a mouse model that can recapitulate FTD-related pathologies. We employed an adeno-associated virus (AAV)-mediated somatic transgenic approach that transduces a gene of interest in the postnatal mouse brain [40, 41]. On postnatal day 0 (p0), mice were administered intracerebroventricular (ICV) microinjections of AAV expressing HA-tagged M719V-CYLD and EGFP (*HA-M719V* mice), HA-CYLD and EGFP (*HA-CYLD* mice), or EGFP (*EGFP* mice) alone, with *EGFP* mice serving as a control while labeling infected neurons (Fig 1.a). Three months after ICV administrations, strong GFP fluorescence was detected throughout the brain including, but not limited to, the PFC, hippocampus, striatum, hippocampus, and thalamus (Fig. 1b,c). As expected with the neuronal promoter human *Synapsin 1* (*hSyn*), AAV-mediated expression specifically occurred in neurons (Fig. 1d), but not in microglia (Fig. 1e). Western blot analysis of CYLD in cortical lysates revealed approximately 3-fold overexpression of WT-CYLD or M719V-CYLD compared to control mice (Fig. 1f,g). The actual extent of overexpression in neurons may be higher since endogenous CYLD is also expressed in non-neuronal cells.

**Figure 1.**
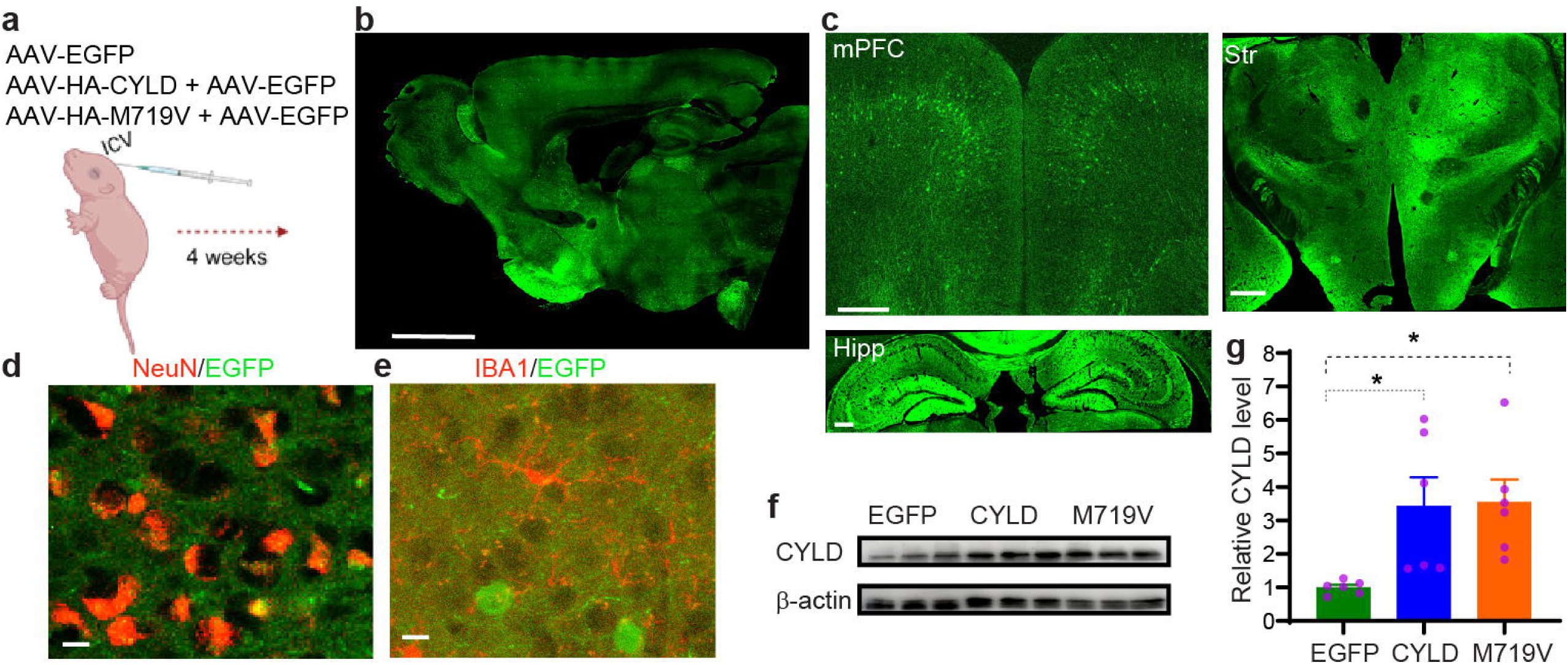
Generation of somatic transgenic mouse models. a. Schematic of intracerebroventricular (ICV) viral (co)injections at postnatal day 0 (p0). AAV2/9-hSyn-EGFP and AAV2/9-hSyn-3xHA-CYLD, AAV2/9-hSyn-EGFP and AAV2/9-hSyn-3xHA-M719V-CYLD, or AAV2/9-hSyn-EGFP alone were injected. b. A sagittal section from a 3-month-old somatic EGFP mouse showing viral spread and expression throughout the mouse brain. Scale bar, 500 μm. c. Coronal sections showing EGFP expression in the mPFC, striatum and hippocampus. Scale bars, 250 μm. d. Co-expression of NeuN and EGFP in *EGFP* mice. Scale bar, 20 μm. e. Lack of co-expression of IBA1 and EGFP in *EGFP* mice. Scale bar, 20 μm. f. Representative blots of CYLD and β-actin on somatic *EGFP*, *HA-CYLD*, and *HA-M719V* mouse cortical lysates. g. Quantification of CYLD protein levels normalized to *EGFP* mice. n = 6 each group. Statistics: One-way ANOVA followed by Tukey’s post hoc tests. *p < 0.05. Data represents mean ± SEM.

### *M719V-CYLD* somatic transgenic mice display early-onset and progressively worsening behavioral deficits

We first tested *EGFP*, *HA-CYLD* and *HA-M719V* somatic transgenic mice for FTD-related behavioral deficits by subjecting them to a behavioral battery at increasing ages (3, 6, and 9 months). In the open field test (OFT) (Fig. 2a), all three mouse groups traveled similar total distances at 3, 6 and 9 months, suggesting unaltered basal locomotor activity (Fig 2b). However, *HA-M719V* and *HA-CYLD* spent more time at the center of the open field arena at 9 but not 3 or 6 months old compared to *EGFP* mice, suggesting age-dependent decrease in anxiety in these groups (Fig 2b).

**Figure 2.**
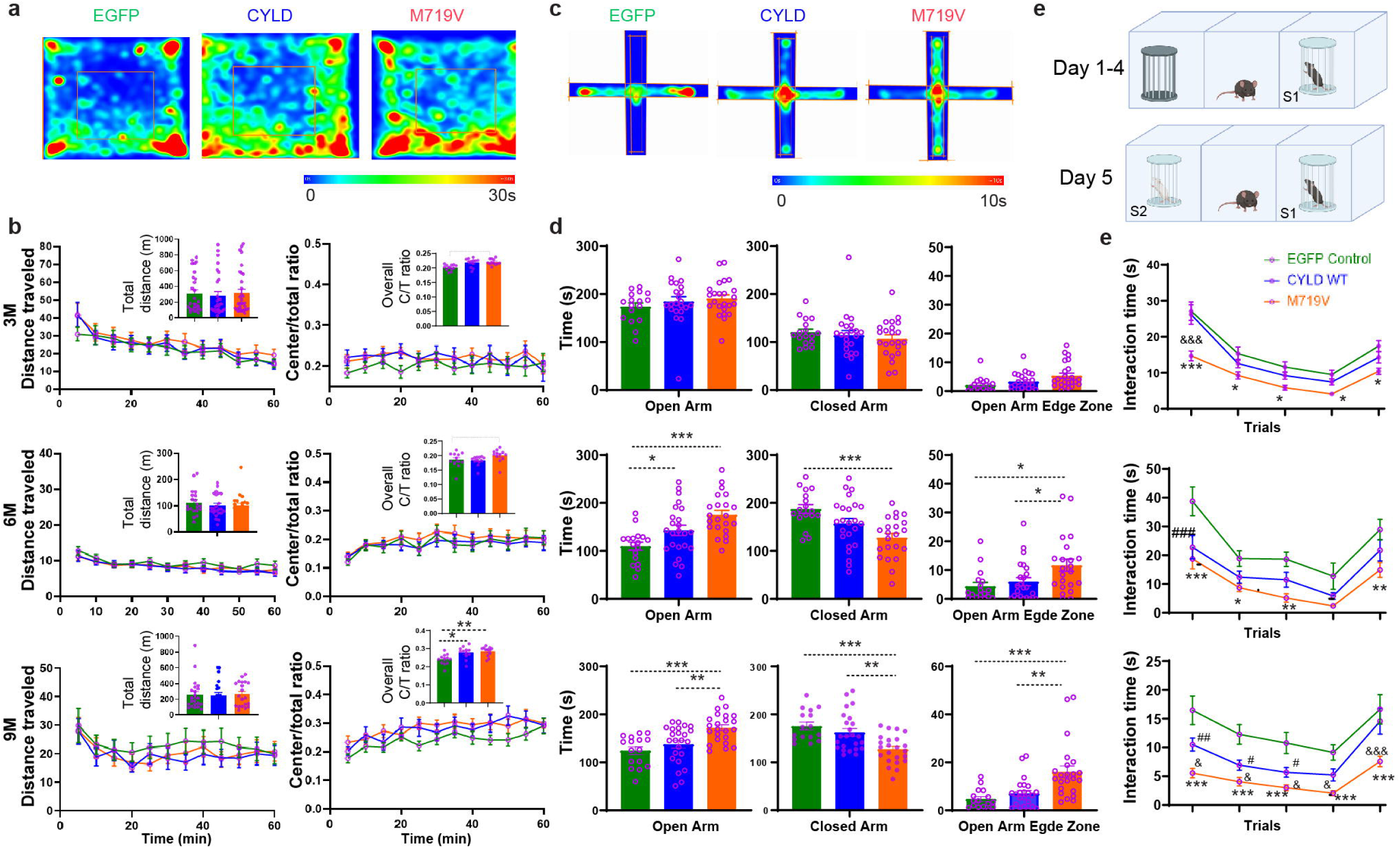
Early onset and progressively worsening anxiety, risk-taking, and social behavioral deficits in somatic *M719V* transgenic mice. a. Representative heatmaps of locomotor activity of 9-month-old *EGFP*, *HA-CYLD* and *HA-M719V* somatic transgenic mice in OFT. b. Time courses of distance traveled and center-to-total distance ratios in 3-, 6-, and 9-month-old *EGFP*, *HA-CYLD* and *HA-M719V* mice in OFT. Insets, total distance traveled and center-to-total distance ratios in 60 min. N = 26 (*EGFP*), 29 (*HA-CYLD*) and 32 (*HA-M719V*) mice at 3 months; n = 20, 20, and 22 at 6 months; and n = 20, 19, and 19 at 9 months. c. Representative heatmaps of time spent in EPM of 9-month-old *EGFP*, *HA-CYLD* and *HA-M719V* mice. d. Time spent on the Open Arm, Closed Arm, and Open Arm Edge Zones in 3-, 6-, and 9-month-old *EGFP*, *HA-CYLD* and *HA-M719V* mice in EPM. N = 17 (*EGFP*), 22 (*HA-CYLD*) and 23 (*HA-M719V*) mice at 3 months; n = 17, 24, and 22 at 6 months; and n = 16, 24, and 23 at 9 months. e. Schematic of a 5-trial three-chamber social behavioral assay. f. Total interaction time with test mice (stranger S1 or novel S2) across 5 consecutive trials. n = 18 (*EGFP*), 21 (*HA-CYLD*), and 20 (*HA-M719V*) at 3 months; n = 15, 16, and 18 at 6 months; and n = 16, 20, and 24 at 9 months. Statistics: One-way or two-way ANOVA followed by Turkey’s post hoc tests. *p < 0.05, **p < 0.01, ***p < 0.001, *M719V* vs. *EGFP*; ^#^p < 0.05, ^##^p < 0.01, ^###^ p < 0.001, *CYLD* vs. *EGFP*; ^&^p < 0.05, ^&&^p < 0.01, ^&&&^p < 0.001, *M719V* vs. *CYLD*.

In the elevated plus maze (EPM) test (Fig. 2c), *HA-M719V* mice spent similar amount of time on open or closed arms as *EGFP* and *HA-CYLD* mice at 3 months of age, but more time on the open arm (and accordingly less time on the closed arm) than *EGFP* mice but not *HA-CYLD* mice at 6 months of age, and more time than both *EGFP* and *HA-CYLD* mice at 9 months of age, consistent with an age-dependent reduction of anxiety. Interestingly, a similar trend of reduced anxiety, albeit to a lesser degree, was also found in *HA-CYLD* mice compared to *EGFP* mice as they spent more time on the open arm compared to *EGFP* mice at both 6 and 9 but not 3 months of age (Fig. 2d), suggesting elevated CYLD activity is responsible for this phenotype. Furthermore, *HA-M719V* mice spent significantly more time on the edges of open arms compared to both *EGFP* and *HA-CYLD* mice at all three ages, suggesting risk-taking-like behaviors in *M719V* mutant mice even at very early stages of disease (Fig. 2d).

Impaired social behaviors are common in FTD especially in behavioral variant of FTD (bvFTD) [1, 2]. In the 5-trial 3-chamber social behavioral test [36, 43] (Fig. 2e), *HA-M719V* mice exhibited marked impairments in both sociability and social novelty compared to *HA-CYLD* and *EGFP* mice that emerged at 3 months and persisted through and worsened at 9 months (Fig. 2f). Again, compared to *EGFP* mice, *HA-CYLD* mice showed less pronounced but significant age-dependent sociability impairments than did *M719V* mice, again indicating elevated CYLD activity is responsible for the observed behavioral phenotype. Social memory appeared normal in both *HA-M719V* and *HA-CYLD* somatic transgenic mice compared to *EGFP* mice at all ages examined.

Empathy loss of is a hallmark feature of bvFTD [1, 2, 44]. To evaluate empathic functions in mice, we employed a distress induced affiliative (DIA) behavioral paradigm (Fig. 3a) that assesses both affective empathy and empathy-driven consolation behaviors [41]. Compared to *EGFP* mice, *HA-M719V* mice displayed strikingly reduced vicarious freezing during the observational fear conditioning (OFC) session of DIA at 3, 6, and 9 months, suggesting impaired social transmission of fear in these mice even at very young ages (Fig. 3b). Observational fear in *HA-CYLD* mice was unchanged at 3 months, modestly decreased compared to *EGFP* mice but remained significantly higher than *HA-M719V* mice at 6 months, but severely reduced to the same levels as *HA-M719V* mice at 9 months (Fig. 3b). Furthermore, *HA-M719V* mice exhibited markedly reduced other-directed affiliative behaviors, measured as duration, number of bouts and latency of allogrooming and body contact bouts towards distressed demonstrator mice, beginning at 3 months and worsening at 6 through 9 months, suggesting age-dependent impairments in consolation behavior (Fig. 3c,d; Supplemental Fig. 1). Similarly, consolation behaviors were also impaired in *HA-CYLD* mice to intermediate levels between *EGFP* and *HA-M719V* mice at the three ages (Fig. 3c,d; Supplemental Fig. 1). Taken together, these results indicate that overexpressing M719V-CYLD, and to a lesser extent the wildtype protein, in neonatal neurons elicit striking bvFTD-associated behavioral impairments that begin at early adult but progressively worsen with aging.

**Figure 3.**
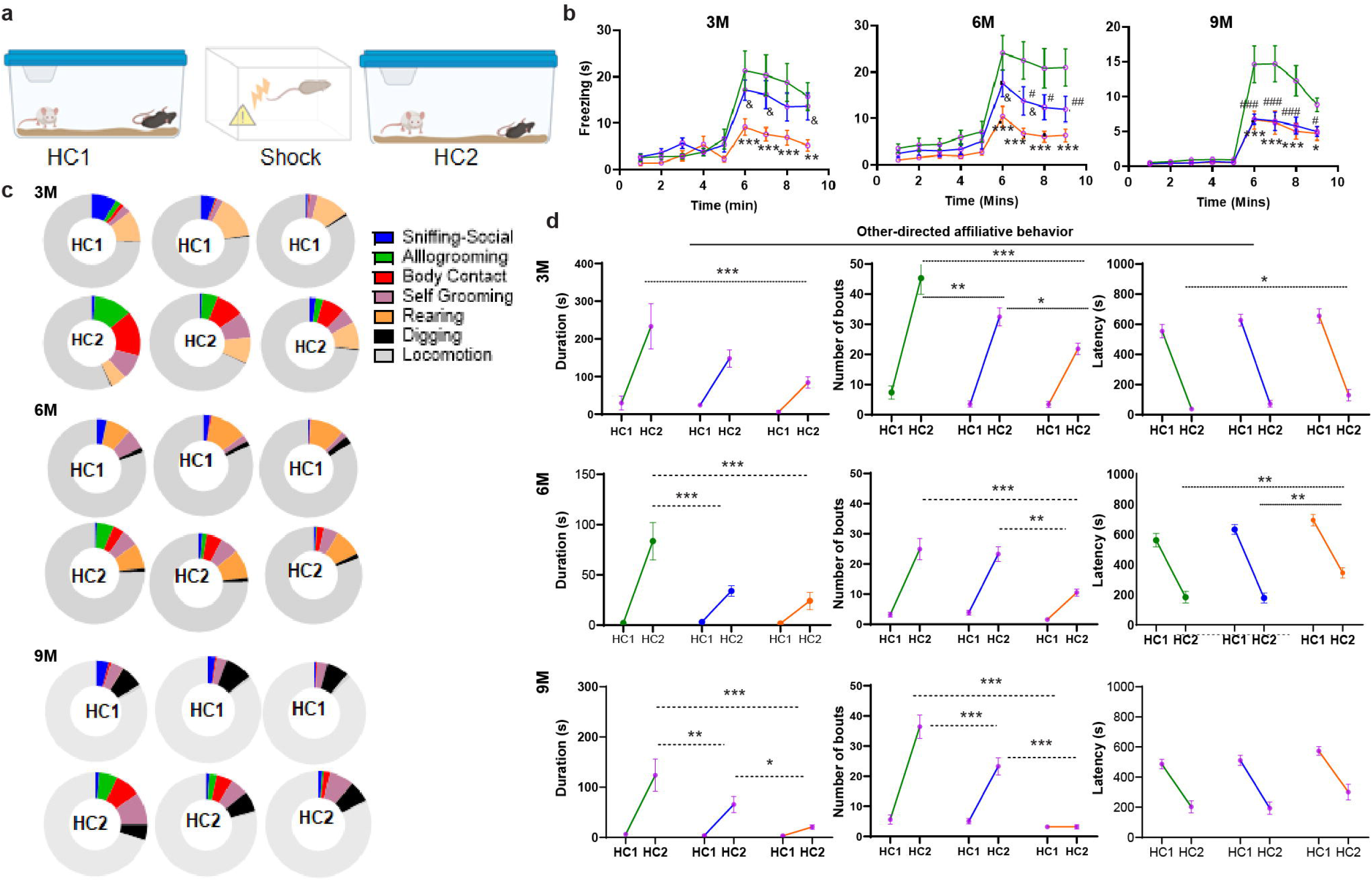
Early onset but progressively blunted empathic functions in *HA-M719V* mice. a. Schematic of DIA paradigm. b. Total freezing time by 3-, 6-, and 9-mopnth-old *EGFP*, *HA-CYLD* or *HA-M719V* observers during OFC. c. Proportions of time 3-, 6-, and 9-mopnth-old *EGFP*, *HA-CYLD* and *HA-M719V* observers spent engaging in different behaviors in HC1 and HC2 during DIA. d. Total duration, number of bouts, and latency of affiliative allogrooming and body contact behaviors by 3-, 6-, and 9-mopnth-old observers toward distressed demonstrators. n = 11 (*EGFP*), 11 (*CYLD)* and 12 (*M719V*) mice at 3 months; n = 16 (*EGFP*), 24 (*CYLD*) and 22 (*M719V*) mice at 6 months; n = 16 (*EGFP*), 23 (*CYLD*) and 21 (*M719V*) mice at 9 months. Statistics: one-way or two-way ANOVA followed by Tukey’s post hoc tests. *p < 0.05, **p < 0.01, ***p < 0.001, *M719V* vs. *EGFP*; ^#^p < 0.05, ^##^p < 0.01, ^###^p < 0.001, *CYLD* vs. *EGFP*; ^&^p < 0.05, ^&&^p < 0.01, ^&&&^p < 0.001, *M719V* vs. *CYLD*.

### *Neuronal M719V-CYLD* overexpression induces profound intrinsic and synaptic deficits in mPFC pyramidal neurons

We next investigated how M719-CYLD affects the neurophysiological properties of prefrontal cortex neurons using slice electrophysiology. To ensure recording and characterizations of individually infected, genetically defined neurons, we administered ICV injections of AAVs encoding EGFP-tagged M719V-CYLD (EGFP-M719V-CYLD), CYLD (EGFP-CYLD), or EGFP on P0 pups, and performed whole-cell patch-clamp recoding from GFP-positive pyramidal neurons on slices prepared from 6- or 9-month-old mice (Fig. 4a). To fit *EGFP*-tagged *CYLD* into the AAV vector, a segment of vector backbone was deleted (please see Methods for more details). These somatic transgenic mice expressing EGFP-M719V-CYLD or EGFP-CYLD showed similar behavioral phenotypes as mice expressing their HA-tagged counterparts at similar ages (Supplemental Figs. 2 and 3).

**Figure 4.**
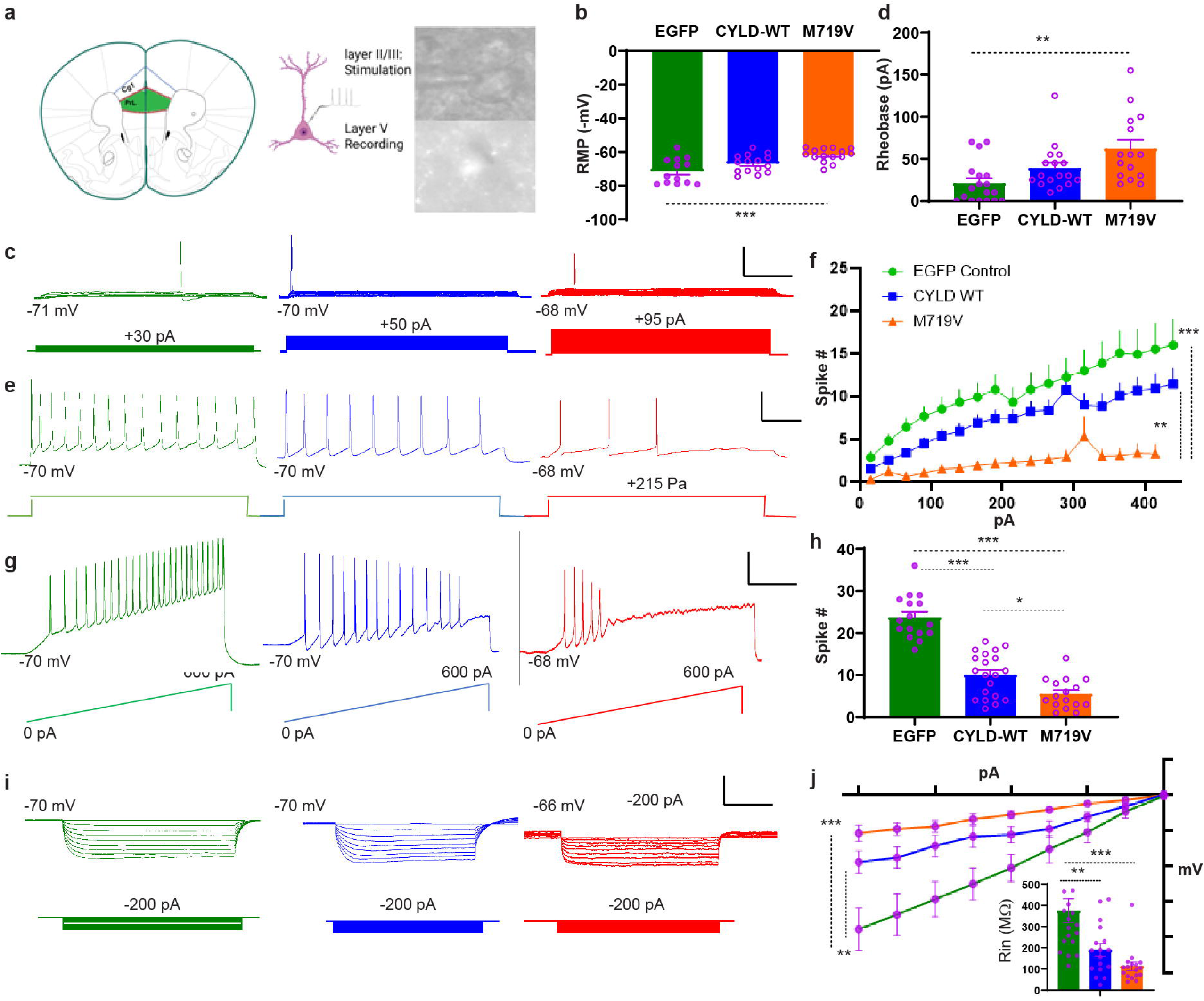
Impaired prefrontal membrane excitability in *EGFP-M719V* somatic transgenic mice. a. Schematic of mPFC recording configuration. Patch-clamp recordings were made on individual layer 5 fluorescent pyramidal neurons in the mPFC expressing EGFP, EGFP-CYLD, or EGFP-M719V-CYLD on slices prepared from 6–9-month-old somatic transgenic mice. b. Summary of RMP in *EGFP*, *EGFP-CYLD* and *EGFP-M719V* mice. n = 13 (*EGFP*), 15 (*EGFP-CYLD*) and 15 (*EGFP-M719V*) cells. c. Representative records showing minimal current injections required to evoke the first spikes in *EGFP*, *EGFP-CYLD* and *EGFP-M719V* neurons. Scale bar: 20 mV, 100 ms. d. Quantification of c. n = 12 (*EGFP*), 17 (*EGFP-CYLD*), and 15 (*EGFP-M719V*) cells. e. Representative membrane potential in response to 215-pA depolarizing current in *EGFP*, *EGFP-CYLD* and *EGFP-M719V* neurons. Scale bar: 20 mV, 100 ms. f. Summary number of spikes vs increasing current injections (500 ms, +25 pA steps) in *EGFP*, *EGFP-CYLD* and *EGFP-M719V* neurons. n = 15 (*EGFP*), 13 (*EGFP-CYLD*), and 14 (*EGFP-M719V*) cells. g. Representative voltage responses to a depolarizing ramp current in *EGFP*, *EGFP-CYLD* and *EGFP-M719V* neurons. Scale bar: 20 mV, 100 ms. h. Quantification of g. Sample sizes: n = 19 (*EGFP*), 17 (*EGFP-CYLD*), and 17 (*EGFP-M719V*) cells. i. Representative voltage responses to a hyperpolarizing step currents in *EGFP*, *EGFP-CYLD* and *EGFP-M719V* cells. Scale bar: 20 mV, 100 ms. j. Summary voltage-current relationships quantified from i. Insets, Summary of input resistance. n = 18 (*EGFP*), 16 (*EGFP-CYLD*), and 16 (*EGFP-M719*) cells. Statistics: *p < 0.05, **p < 0.01, ***p < 0.001; One-way or two-way ANOVA with Tukey’s post hoc tests.

We first assessed the intrinsic membrane properties of layer 5 pyramidal neurons (L5PNs) in the mPFC under current clamp (Fig. 4). At 6-9 months of age, the resting membrane potential (RMP) was significantly depolarized in mPFC L5PNs expressing EGFP-M719V-CYLD than EGFP-expressing neurons, suggesting compromised neuronal health in mutant neurons (Fig. 4b). Compared to both EGFP and EGFP-CYLD neurons, *EGFP-M719V-CYLD* neurons displayed increased rheobase (minimal current required to evoke an action potential) (Fig. 4c,d), reduced firing rate in response to either step (Fig. 4e,f) or ramp (Fig. 4,h) depolarizing currents, and decreased input resistance (a measure of membrane leakage; Fig. 4i,j). These data demonstrate loss of neuronal excitability due to leaky cell membrane in mutant neurons.

We next investigated the synaptic properties of mPFC L5PNs (Fig. 5). We recorded AMPA receptor-mediated miniature excitatory postsynaptic currents (mEPSCs) under voltage-clamp (Vh = −70 mV) in the presence of TTX (to block voltage-gated Na+ currents) and picrotoxin (to block GABA_A_ receptor-mediated inhibitory currents). The amplitude of mEPSCs was similar among all groups but the frequency of mEPSC was significantly reduced in *EGFP-M719V-CYLD* neurons compared to *EGFP* neurons (Fig. 5a,b), suggesting a loss of functional synapses and/or a decrease in the presynaptic glutamate release probability. Next, we measured the AMPA/NMDA ratio, an index for synaptic efficacy measured in a kinetics-based method (Fig. 5c) [42]. AMPA/NMDA ratio was significantly reduced in *EGFP-M719V-CYLD* neurons compared to both *EGFP-CYLD* and *EGFP* neurons, consisting with a reduced synaptic strength in mutant mPFC pyramidal neurons.

**Figure 5.**
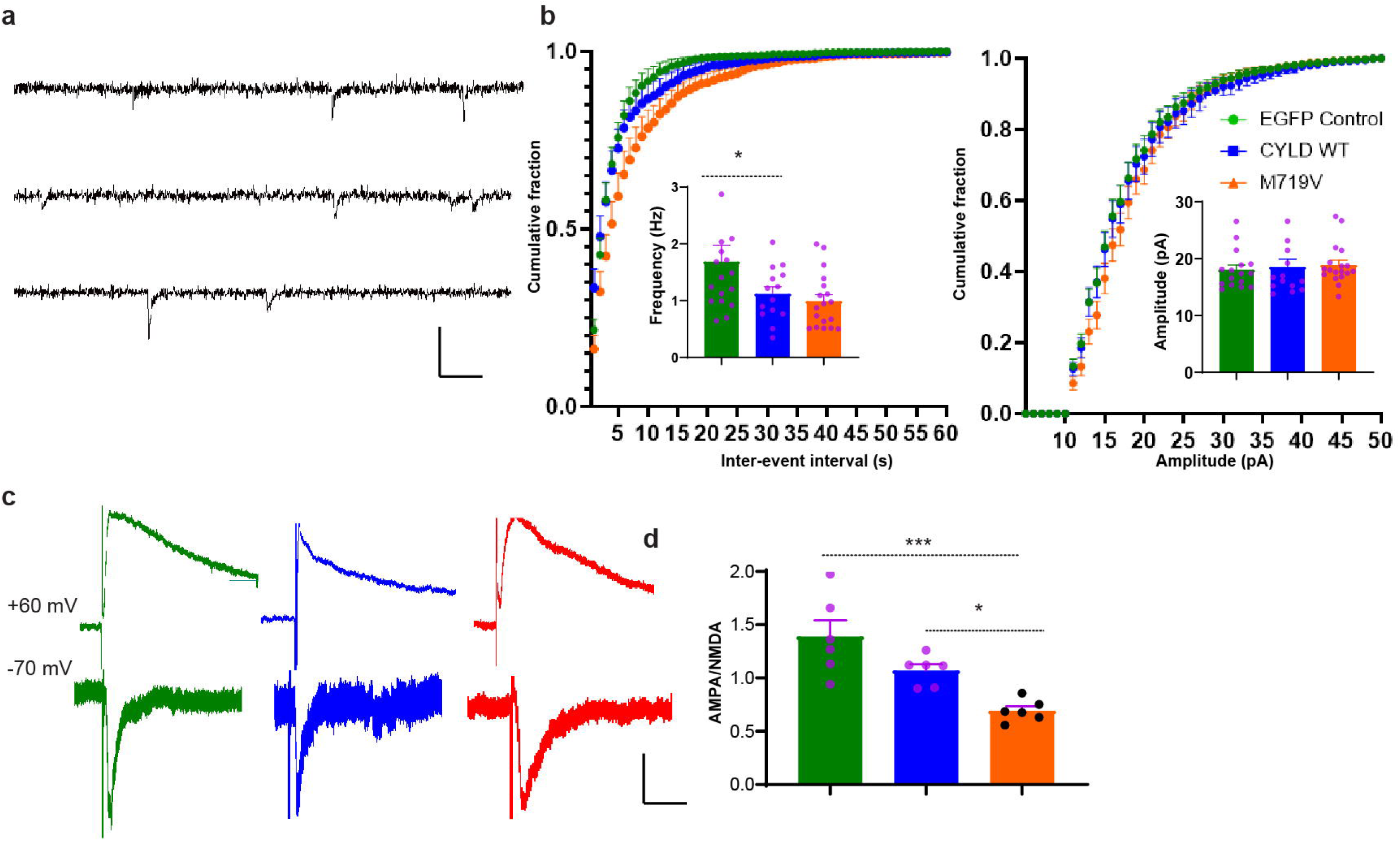
Reduced synaptic efficacy in *EGFP-M719V* somatic transgenic mice. a. Representative mEPSC records from *EGFP*, *EGFP-CYLD* and *EGFP-M719V* neurons. Scale bar, 40 pA, 2 sec, b. Cumulative probabilities of interevent interval (left) and amplitude (right) distributions of mEPSCs. Inserts, mean ± SEM of mEPSC frequency and amplitude. n = 16 (*EGFP*), 14 (*EGFP-CYLD*), and 18 (*EGFP-M719V*) cells. c. Example EPSC recordings at −70 mV (to record AMPAR-mediated responses) and +60 mV (to record both NMDAR and AMPAR-mediated responses). Scale bar, 40 pA, 20 ms. d. Summary AMPA/NMDA ratio, measured as the peak AMPAR component at −70 mV divided by the amplitude of the NMDAR component 80 ms after stimulation at +60 mV. n = 6 (*EGFP*), 6 (*EGFP-CYLD*), and 6 (*EGFP-M719V*) cells. Statistics: *p < 0.05, ***p < 0.001; One-way ANOVA with Tukey’s post hoc tests.

Given the early onset of behavioral symptoms in *M719V* somatic transgenic and to a lesser extent *CYLD* somatic transgenic mice, we hypothesize that neuronal M719V-CYLD exerts its functionally toxic effects more acutely. To test this hypothesis, we stereotaxically injected *EGFP-M719V-CYLD*, *EGFP-CYLD*, or *EGFP* AAVs locally into the mPFC of young adult (2-month) mouse brain and performed slice electrophysiology 1-2 months later (Supplemental Figs. 4 and 5). We observed nearly identical results as above: EGFP-M719V-CYLD-expressing L5PNs, and to a lesser extent EGFP-CYLD-expressing L5PNs, displayed more depolarized RMPs (Supplemental Fig. 4), reduced membrane excitability (Supplemental Fig. 4), and decreased synaptic strength (Supplemental Fig. 5) compared to EGFP-expressing L5PNs. These data suggest that the neuronal overexpression of CYLD elicits rapid functional remodeling of adult PFC neurons and FTD-associated M719V-CYLD exhibits a gain-of-function property.

### Mild neuropathologies in *M719V* somatic transgenic mice

Patient brains of FTD, including *CYLD*-FTD, are characterized by neuronal loss, microgliosis, and TDP-43 pathologies [2, 3]. Thus, we examined our somatic transgenic mouse models for these FTD-associated pathologies in the mPFC at 12 months of age. The number of NeuN-positive neurons in the mPFC was statistically similar among *HA-CYLD*, *HA-M719V*, and *EGFP* mouse groups (Fig. 6a,b), suggesting lack of obvious neurodegeneration. Numbers of IBA1 (Ionized calcium-binding adaptor molecule 1, a marker of microglia)-positive cells and CD11b (Integrin alpha M/ITGAM, a marker of activated microglia) levels were both statistically similar among genotypes (Fig. 6e,f), suggesting neither microglial proliferation nor activation was significantly altered in *M719V-CYLD* or *CYLD* mice. Finally, we observed a significant decrease of p62 (an autophagy substrate and a proteostasis marker)-positive clusters in both *HA-CYLD* and *HA-M719V* mice compared to *EGFP* mice (Fig. 6c,d), indicating enhanced clearance of autophagic cargos. Together, these data suggest that neuronal overexpression of M719V-CYLD does not induce overt FTD-related neuronal cell loss of microgliosis even at 12 months of age.

**Figure 6.**
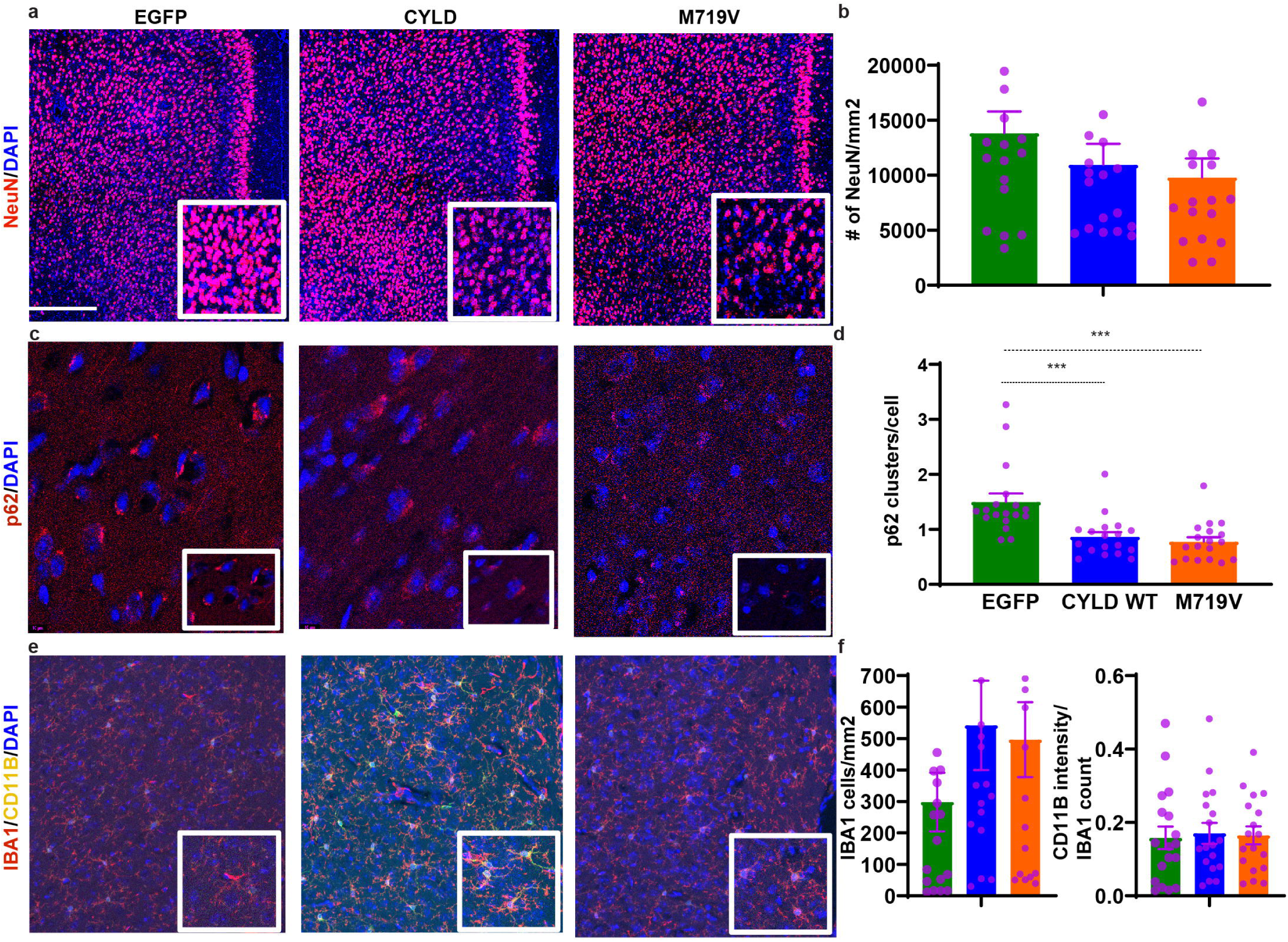
Neuropathological assessment of 12-month-old *HA-M719V* mice. a. Representative NeuN immunostaining from 12-month-old *EGFP*, *HA-CYLD*, and *HA-M719V* mice. Nuclei are labeled with DAPI. Scale bar, 100 μm. b. Quantification NeuN cell counts from a. c. Representative p62 immunostaining from 12-month-old *EGFP*, *HA-CYLD*, and *HA-M719V* mice. Nuclei are labeled with DAPI. Scale bar, 100 μm. d. Quantification of p62 clusters per cell from e. e. Representative co-immunostaining of IBA1 and CD11b from 12-month-old *EGFP*, *HA-CYLD*, and *HA-M719V* mice. Nuclei are labeled with DAPI. Scale bar, 100 μm. f. Quantification IBA1-positive cell counts (per mm2) and CD11b intensity per IBA1 cell) from g. n = 18 sections from 6 mice in each group. Statistics: *p < 0.05, **p < 0.01, ***p < 0.001; One-way ANOVA with Tukey’s post hoc tests.

### Dysregulated Akt-mTOR-autophagy axis in *M719V* somatic transgenic mice

Finally, we explored potential pathogenic mechanisms of *CYLD*-FTD. We previously showed that CYLD, in a DUB enzyme activity-dependent manner, inhibits Akt/mTOR signaling and stimulates induction of neuronal autophagy [36]. Because the *M719V* mutation has been shown to exhibit an increased DUB enzyme activity [3], we tested the hypothesis that M719V-CYLD dysregulates the Akt-mTOR-autophagy axis in mutant mice. Levels of total and phosphorylated Akt (T308) were equally significantly reduced in cortical lysates prepared from 12-month-old *HA-CYLD* and *HA-M719V* mice compared to *EGFP* mice (Fig. 7a,b). Levels of total and phosphorylated mTOR (a master inhibitor of autophagy) were modestly reduced in *HA-CYLD* and *HA-M719V* mouse cortical lysates, but most of the differences did not reach a significant level (Fig. 7a,c). Consistently with above IHC data, levels of the autophagy adaptor p62 were similarly and significantly reduced in *HA-CYLD* and *HA-M719V* cortical lysates (Fig. 7a,d). Finally, both LC3-I and LC3-II (the lipidated form of LC3-I and a reliable marker for autophagosomes) were upregulated in *HA-CYLD* and *HA-M719V* mouse cortex, with LC3-I in *HA-M719V* mice showing the most striking increase (Fig. 7a,e). Interestingly, however, the LC3-II/I ratio was unaltered or slightly decreased in *HA-CYLD* and *HA-M719V* mice. These data reveal complex dysregulations of the Akt-mTOR-autophagy axis, resulting in a net increase of autophagy activity in *M719V* mice and to a lesser extent in *CYLD* mice.

**Figure 7.**
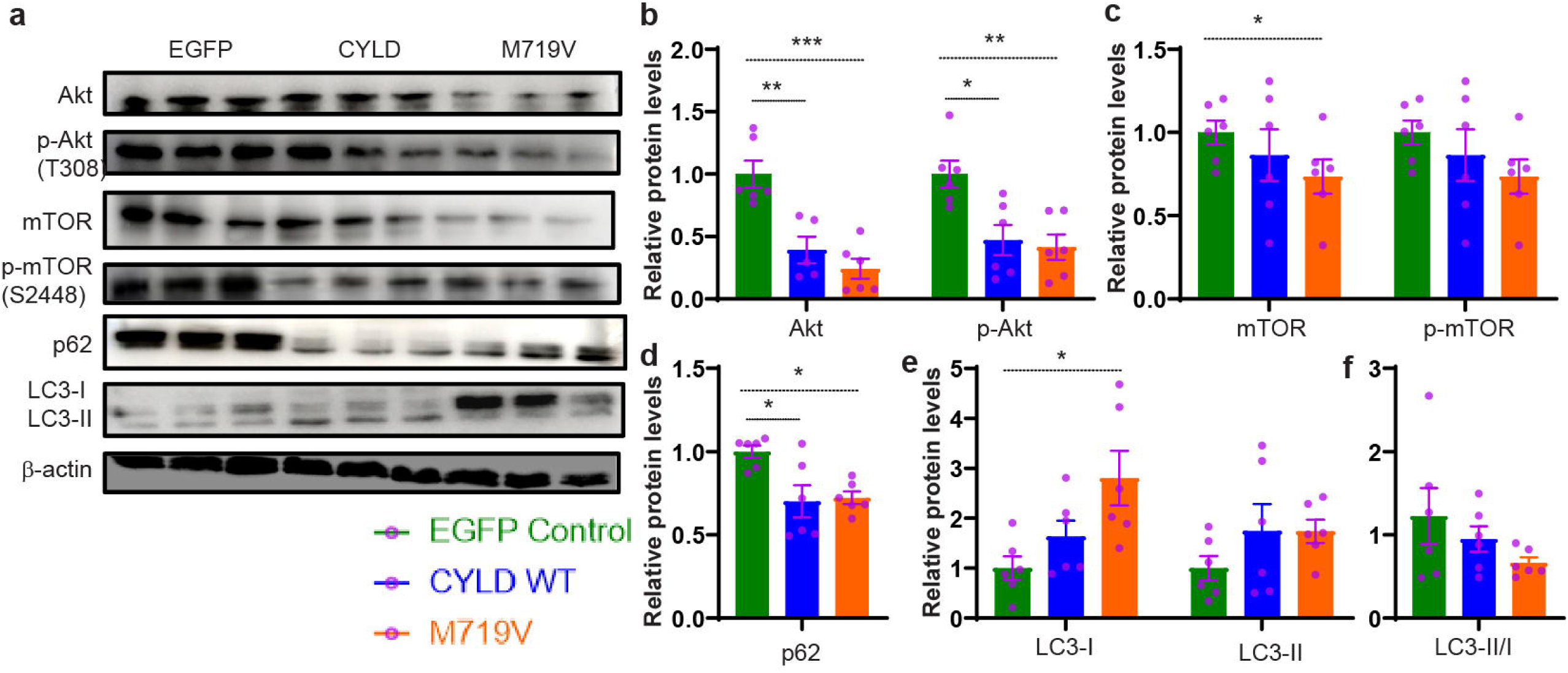
Altered Akt-mTOR-autophagy signaling axis in 12-month-old *HA-M719V* mice. a. Representative blots for indicated proteins from cortical lysates prepared from 12-month-old *EGFP*, *HA-CYLD*, and *HA-M719V* mice. b-f. Densitometric quantification of indicated proteins. n = 6 mice in each group. *p < 0.05, **p < 0.01, ***p < 0.001; One-way ANOVA with Tukey’s post hoc tests.

## Discussion

In this study, we establish a somatic transgenic mouse model of *CYLD*-FTD and investigate the molecular, neurophysiological and behavioral pathologies associated with the M719V mutation. We show that neonatal expression of M719V-CYLD in postnatal neurons elicits profound and early (as early as 3 months) neurophysiological and behavioral impairments yet no obvious neurodegeneration and molecular pathologies as late as 12 months of age. Furthermore, overexpressing even the wild-type CYLD has similar, albeit less robust molecular, physiological and behavioral effects, indicating that CYLD activation is responsible for the observed behavioral phenotypes as the M719V mutation enhances CYLD activity. Finally, we found complex dysregulations of the Akt-mTOR-autophagy signaling resulting in an elevated autophagy activity in mutant mouse cortex. These results establish the first animal model of *CYLD*-FTD and reveal that CYLD activation can lead to FTD-related behavioral deficits.

Given the autosomal dominant nature of *CYLD*-FTD in patients [3], the lack of apparent neuronal loss and microgliosis in young *M719V* mouse brain or even at 12 months suggests that profound functional impairments at neurophysiological and behavioral levels are early disease phenotypes in *CYLD*-FTD and possibly other forms of FTD as well. Indeed, despite a more depolarized RMP, consistent with an unhealthy state of mutant neurons, prefrontal M719V-CYLD-expressing neurons exhibit strikingly decreased membrane excitability (which might be counterintuitive but not surprising), characterized by an increased rheobase, a reduced input resistance, and a lower firing rate. In parallel, synaptic strength in mutant prefrontal neurons is also significantly diminished, characterized by a lower mEPSC frequency and a decreased NMDA/AMPA ratio. These synaptic impairments are consistent with M719V-CLYD being a gain of function mutant of the wild-type CYLD that generally inhibits synaptic strength and plasticity [31, 36]. These intrinsic and synaptic impairments are correlated with deficient inhibitory control, affective and social behaviors, consistent with the importance of CYLD in maintaining a repertoire of behaviors [36, 38, 39]. Importantly, much of the physiological and behavioral deficits in *M719V* mice emerge as early as 3 months, supporting functional alterations at synaptic, cellular and circuit levels rather than structural degeneration in mutant mice. Consistent with a more acute role of neuronal CYLD in the postnatal CNS, introducing M719V-CYLD into young adult mouse brain leads to rapid and similar intrinsic and synaptic impairments several weeks later. Hallmark behavioral symptoms of FTD, including changes in personality, loss of empathy, and disinhibition often present at early stages, followed by general memory and cognitive deteriorations at later stages [1]. Thus, early prodromal synaptic and circuit dysfunctions mediating behavior impairments may precede massive neuronal cell loss and disability, but the underling mechanisms are not well known. Our study identifies CYLD/M719V-CYLD-dependent circuit remodeling as such a prodromal mechanism that may contribute to bvFTD onset, progression and diagnosis.

A notable feature of our study is that merely overexpressing the wild-type CYLD protein already has similar, albeit lesser effects on behavior, neurophysiology, and molecular and cellular pathologies than M719V-CYLD overexpression. This may not be surprising because M719V-CYLD possesses an increased DUB enzyme activity [3], thus is expected to behave as a gain-of-function mutant. The DUB enzyme activity is required for much of CYLD’s function in regulating K63 ubiquitinatipon, PSD-95 clustering, and dendritic spine remodeling in neurons [31]. Thus, it is possible that CYLD variant-mediated FTD pathogenesis may rely on DUB enzymatic activity in a dose dependent manner. Nevertheless, potential involvements of DUB-independent mechanisms, such as alterations of CYLD interactions with other FTD-related risk proteins (e.g. p62, TBK1, and optineurin), cannot be excluded and require further investigation.

Autophagy dysregulation has emerged as a major pathogenic mechanism in FTD/ALS [4, 8, 9]. At least 10 of ∼15 FTD genes (e.g. SQSTM1, OPTN, TBK1, and c9orf72) encode direct regulators of the autophagy cycle, and their mutations, mechanistically, impair autophagy functions[4, 8, 45]. Abundant studies have also shown that autophagy is impaired in many FTD mouse models and patient brains[8, 9]. However, it remains enigmatic whether autophagy is hyperactive or compromised in FTD/ALS because either promoting or inhibiting autophagy flux can alleviate neurodegeneration in models of FTD/ALS caused by different genetic mutations[46, 47]. Here, we provide evidence that *CYLD*-FTD is associated with a paradoxically dysregulated autophagy system, characterized by an upregulated autophagy machinery yet unhanged LC3I-II conversion, suggesting an effectively compromised lysosome degradation mechanism. It remains to be determined how this paradoxical autophagy dysregulations contributes to behavioral and physiological impairments in mutant mice. In summary, our results uncover important roles of neuronal CYLD in brain integrity, function and behaviors and establish a unique animal model to investigate pathogenic mechanisms of FTD at molecular, cellular, synaptic and circuit levels.

## ACKNOWLEDGEMENTS

We thank members of the Yao laboratory for critiques and comments. This work was supported by NIH Grants R56NS122351, R21NS125845, and RF1AG082478 (to W.-D.Y. and F.-B.G.).

