## Supplemental figures for "Elevating Neuronal *CYLD* Causes Frontotemporal Dementia (FTD)-Relevant Behavioral and Physiological Deficits"

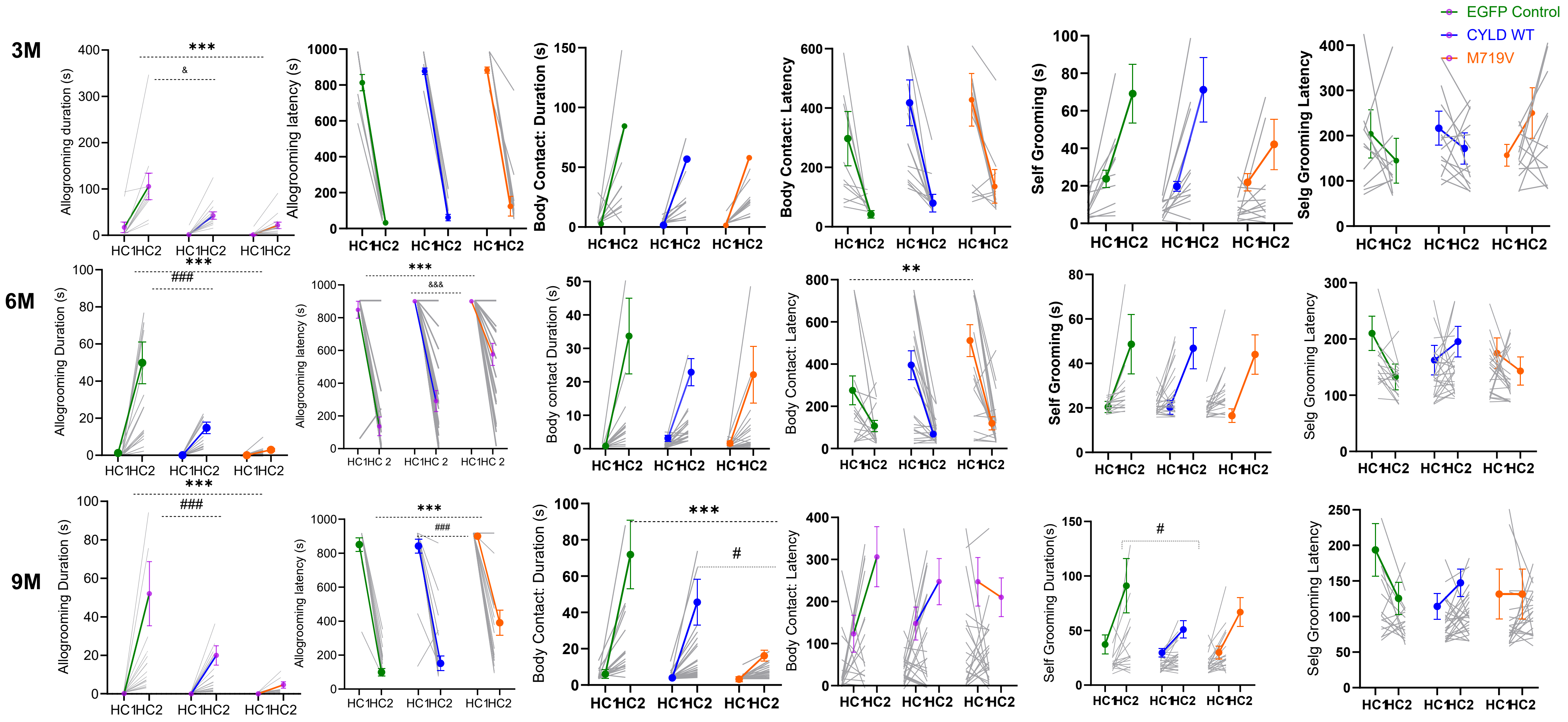


**Supplemental Figure 1**

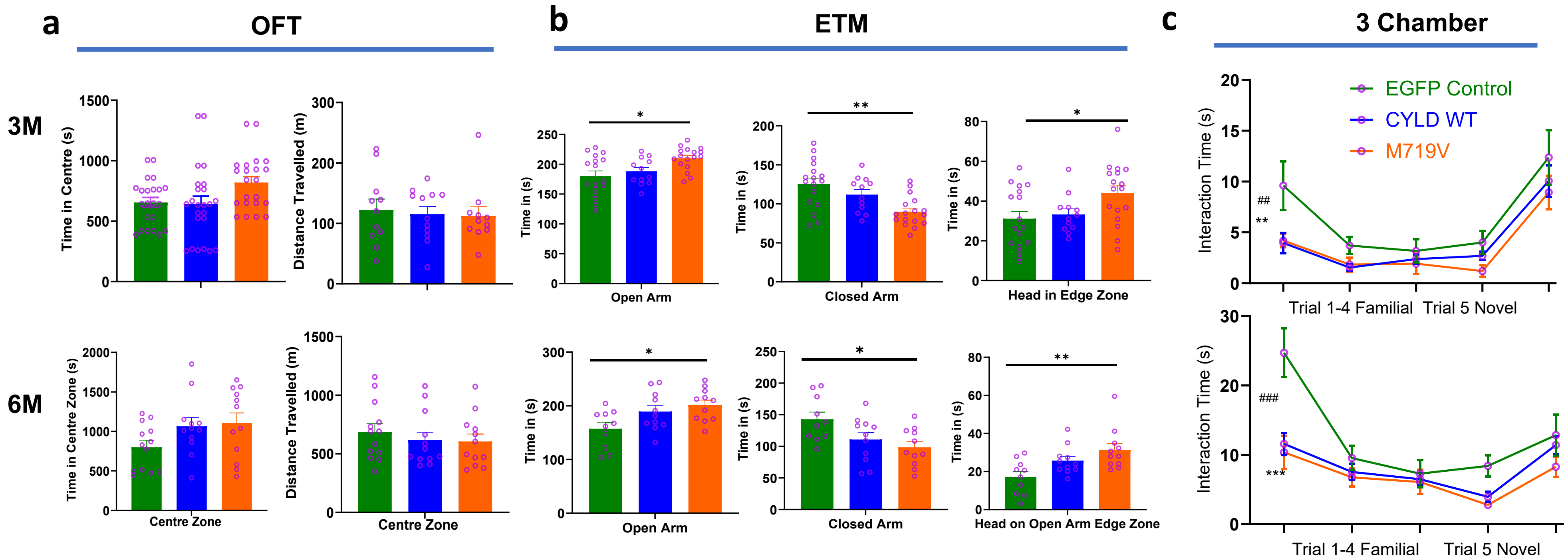


Supplemental Figure 2

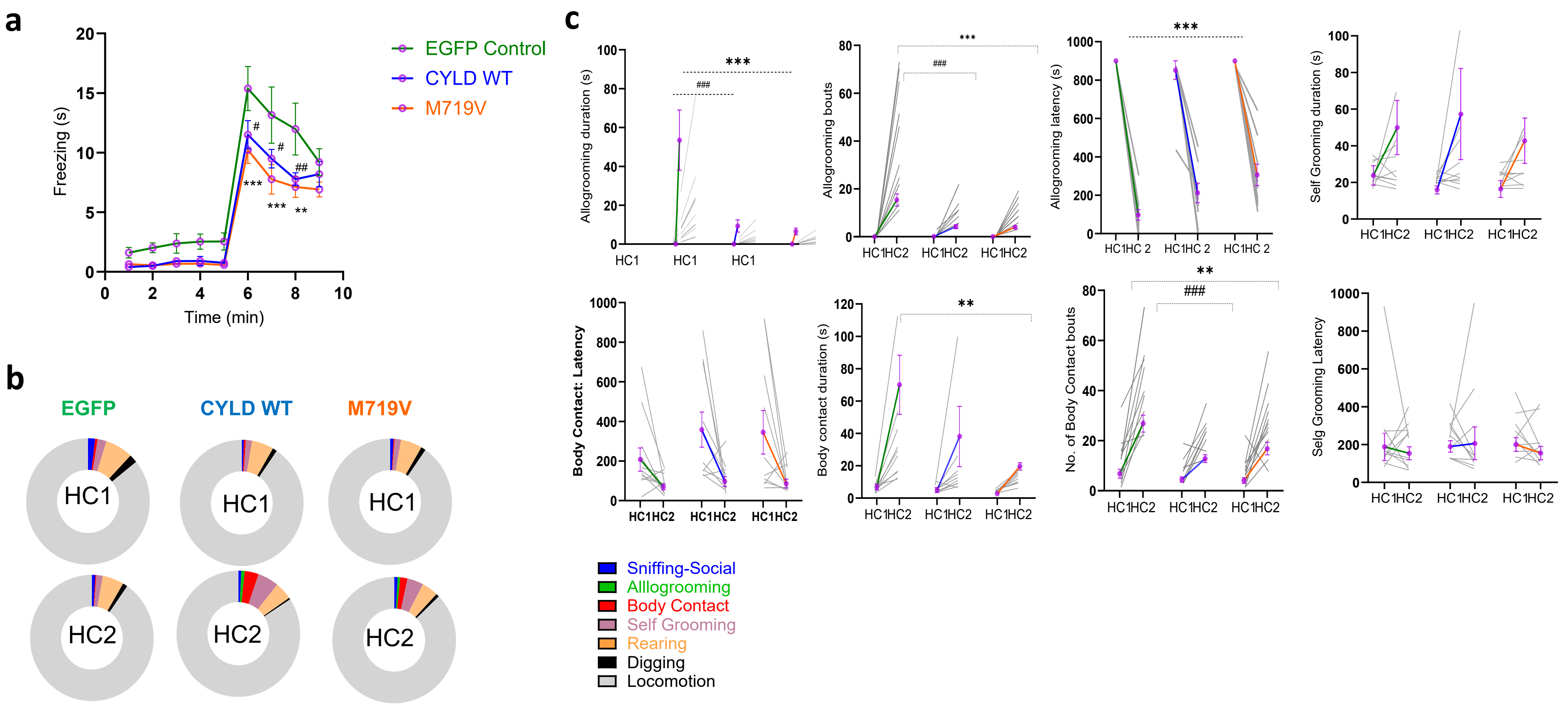


**Supplemental Figure 3**

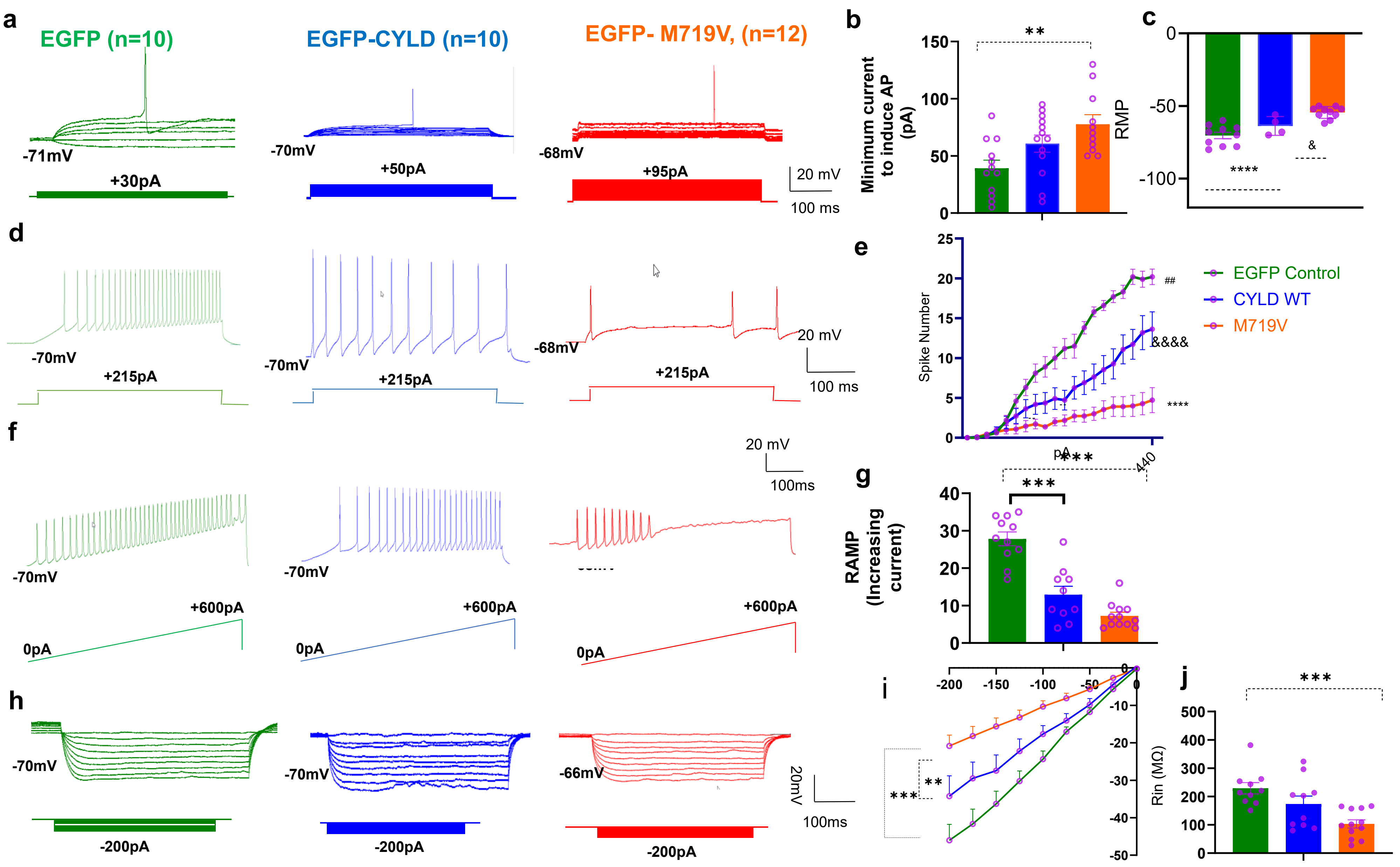


Supplemental Figure 4

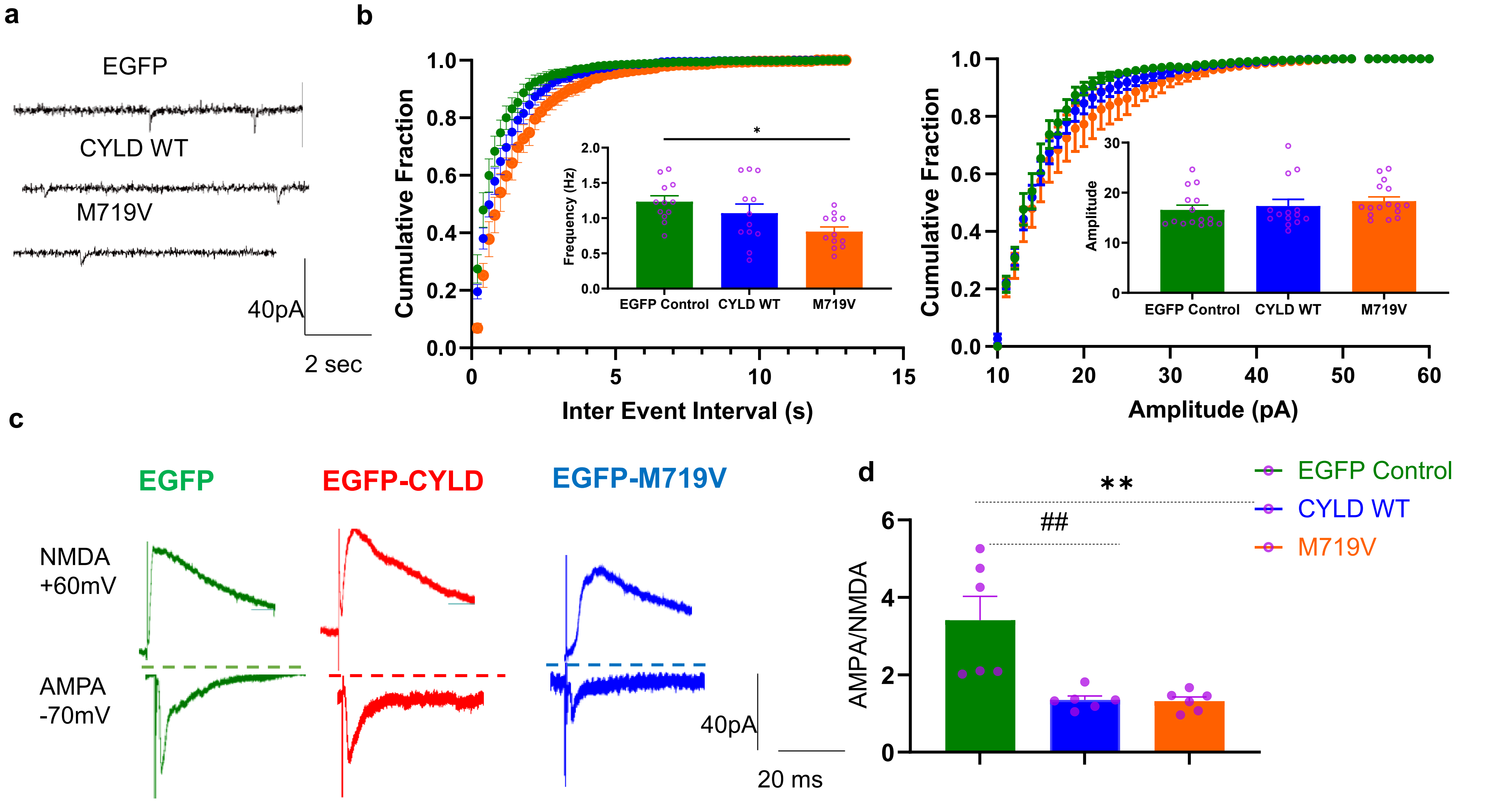


**Supplemental Figure 5**
