## Supplemental figure legends for "Elevating Neuronal *CYLD* Causes Frontotemporal Dementia (FTD)-Relevant Behavioral and Physiological Deficits"

**Figure S1**. Additional behavioral analysis of *EGFP*, *HA-CYLD* and *HA-M719V* somatic transgenic mice in the DIA paradigm. Allogrooming duration and latency, body contact duration and latency, and self-grooming duration and latency of *EGFP*, *HA-CYLD* and *HA-M719V* mice at 3-, 6-, and 9-month-old are shown. n = 10 (*EGFP*), 13 (*CYLD*), and 12 (*M719V*) at 3 months; n = 16, 18, and 22 at 6 months; n = 16, 23, and 21 at 9 months. Statistics: *$p < 0.05$, **$p < 0.01$, ***$p < 0.001$; One-way or two-way ANOVA followed by Tukey's post hoc tests. Data represents mean ± SEM.

**Figure S2**. Behavioral analysis of *EGFP*, *EGFP-CYLD* and *EGFP-M719V* somatic transgenic mice in the open field, elevated plus maze, and three-chamber assays.

a. Total distance traveled and time spent in center zone in 3- and 6-month-old *EGFP*, *EGFP-CYLD* and *EGFP-M719V* mice in OFT. n = 11 (*EGFP*), 11 (*CYLD*), and 11 (*M719V*) at 3 months; n = 12, 12, and 13 at 6 months.
b. Time spent on the Open Arm, Closed Arm, and Open Arm Edge Zones in 3- and 6-month-old *EGFP*, *EGFP-CYLD* and *EGFP-M719V* mice in EPM. n = 17 (*EGFP*), 12 (*CYLD*), and 17 (*M719V*) at 3 months; n = 10, 11, and 11 at 6 months.
c. Total interaction time with test mice across 5 consecutive trials. n = 17 (*EGFP*), 12 (*CYLD-WT*), and 17 (*M719V*) at 3 months; n = 10, 11, and 11 at 6 months.

   Statistics: *$p < 0.05$, **$p < 0.01$, ***$p < 0.001$; One-way or two-way ANOVA followed by Tukey's post hoc tests. Data represents mean ± SEM.

**Figure S3**. Behavioral analysis of 6-month-old *EGFP*, *EGFP-CYLD* and *EGFP-M719V* somatic transgenic mice in the DIA assay.

a. Total freezing time by 6-month-old *EGFP*, *EGFP-CYLD* and *EGFP-M719V* somatic transgenic observers during OFC.
b. Proportions of time 6-mopnth-old observers spent engaging in different behaviors in DIA tests.
c. Total duration, number of bouts, and latency of affiliative behaviors by 6-mopnth-old observers toward distressed demonstrators.

n = 11 (*EGFP*), 10 (*CYLD*), and 10 (*M719V*). Statistics: *p < 0.05, **p < 0.01, ***p < 0.001; One-way or two-way ANOVA followed by Tukey's post hoc tests. Data represents mean ± SEM.

**Figure S4.** Altered mPFC intrinsic membrane properties in acutely infected *EGFP-M719V* neurons.

a. Representative records showing minimal current injections required to evoke the first spikes in layer 5 *EGFP*, *EGFP-CYLD* and *EGFP-M719V* pyramidal neurons in the mPFC. Scale bar: 20 mV, 100 ms. Slices were prepared from 3–4-month-old mice that had stereotaxic viral injections into the mPFC 1-2 month prior (at 2-month old).
b. Quantification of a. n = 10 (*EGFP*), 10 (*CYLD*), and 12 (*M719V*) cells.
c. Summary of RMP in *EGFP*, *EGFP-CYLD* and *EGFP-M719V* neurons. n = 10 (*EGFP*), 10 (*CYLD*), and 12 (*M719V*) cells.
d. Representative membrane potential in response to 215-pA depolarizing current in *EGFP*, *EGFP-CYLD* and *EGFP-M719V* neurons. Scale bar: 20 mV, 100 ms.
e. Number of spikes vs increasing current injections in *EGFP*, *EGFP-CYLD* and *EGFP-M719V* neurons. n = 10 (*EGFP*), 10 (*CYLD*), and 12 (*M719V*) cells.

f. Representative voltage responses in response to a depolarizing ramp current in *EGFP*, *EGFP-CYLD* and *EGFP-M719V* neurons. Scale bar: 20 mV, 100 ms.
g. Quantification of f. n = 12 (*EGFP*), 12 (*CYLD*), and 12 (*M719V*) cells.
h. Representative voltage responses in response to a hyperpolarizing step current in *EGFP*, *EGFP-CYLD* and *EGFP-M719V* neurons. Scale bar: 20 mV, 100 ms.
i. Voltage-current relationships quantified from h. n = 10 (*EGFP*), 10 (*CYLD*), and 12 (*M719V*) cells.
j. Summary of input resistance from i.

Statistics: *$p < 0.05$, **$p < 0.01$, ***$p < 0.001$; One-way or two-way ANOVA followed by Tukey's post hoc tests. Data represents mean ± SEM.

**Figure S5.** Reduced synaptic efficacy in acutely infected *EGFP-M719V* neurons.

a. Representative mEPSC records from *EGFP*, *EGFP-CYLD* and *EGFP-M719V* neurons in stereotaxically injected mice. Scale bar, 40 pA and 2 s.
b. Cumulative probabilities of interevent interval and amplitude distributions of mEPSCs. Inserts, mean ± SEM of mEPSC frequency and amplitude.
c. Example EPSC traces at -70 mV and +60 mV. Scale bar, 40 pA and 20 ms.
d. Summary AMPA/NMDA ratio.

n = 11 (*EGFP*), 12 (*EGFP-CYLD*) and 12 (*EGFP-M719V*) cells. *$p < 0.05$, **$p < 0.01$, ***$p < 0.001$; One-way ANOVA followed by Tukey's post hoc tests. Data represents mean ± SEM.
